# The HMD domain of the PAF complex primes Rad6-Bre1 E3 ligase complexes for H2B ubiquitination

**DOI:** 10.64898/2026.03.01.708808

**Authors:** Ammarah Tariq, Shin Ohsawa, Riccardo Zenezini Chiozzi, Panagiotis Patsis, Charlie Williams, Alessandro Stirpe, Thomas A. Clarke, Konstantinos Thalassinos, Marc Bühler, Thomas Schalch

**Affiliations:** Leicester Institute for Structural and Chemical Biology, Division of Molecular and Cell Biology, University of Leicester, United Kingdom; Friedrich Miescher Institute for Biomedical Research, Fabrikstrasse 24, Basel, Switzerland; Institute of Structural and Molecular Biology, University College London; University College London Mass Spectrometry Science Technology Platform, Division of Biosciences, University College London, London, UK; Institute of Structural and Molecular Biology, Birkbeck College London; Department of Molecular Biology, Faculty of Science, Science III, University of Geneva, CH-1211 Geneva 4, Switzerland; Department of Microbiology, Cornell University, Wing Hall, 123 Wing Drive, Ithaca, New York 14853, USA; School of Biological Sciences, University of East Anglia, Norwich, United Kingdom; University of Basel, Petersplatz 10, Basel, Switzerland

**Keywords:** heterochromatin, euchromatin, E3 ligase, ubiquitin, posttranslational modifications, histones, protein complex, *S*. *pombe*

## Abstract

Mono-ubiquitination of histone H2B (H2Bub) is deposited by Bre1-type E3 ubiquitin ligase complexes during transcription elongation and is critical for chromatin organization, DNA repair, and transcription regulation. In *S. pombe*, this activity is carried out by HULC, a complex of the Bre1 homologs Brl1 and Brl2 with Rhp6 and Shf1. While H2Bub deposition depends on recruitment of HULC to RNA Pol II by the Paf1 complex (PAF1C), the molecular basis for how PAF1C activates HULC has remained unclear. Using AlphaFold modelling, biochemical reconstitution, and functional assays, we define the architecture of HULC as a flexible 1:1:1:1 assembly in which Shf1 stabilizes an elongated coiled-coil hairpin that brings the RING and Rad6-binding domains into proximity. However, for full stimulation of HULC activity, we find that the HMD domain of the PAF1C subunit Prf1 stimulates ubiquitin transfer through a RING-binding region (RBR) that repositions the RING domains adjacent to Rhp6 in a catalytically competent configuration. These findings reveal an activation mechanism in which transcription-associated Prf1 primes HULC for ubiquitin transfer through the Prf1RBR, revealing a critical regulatory interface for H2B ubiquitination.

## Introduction

Monoubiquitination of histone H2B (H2Bub) is found at levels of about 1-1.5% in mouse cells and is highly conserved throughout eukaryotes ^1^. H2Bub is deposited by the ubiquitin-conjugating enzyme Rad6 ^2^ in a complex with the E3 ligase Bre1 in *S. cerevisiae* or a complex of RNF20 and RNF40 in mammalian organisms ^3–6^. In addition, a less well-conserved subunit has been identified as part of the H2Bub ubiquitin ligase complexes in various species ^7,8^. In *S. cerevisiae*, this is Lge1, which has been shown to induce condensate formation ^9^.

H2Bub is tightly associated with transcription elongation, maintenance of chromatin organization, and stimulation of H3K79 and H3K4 methylation ^10–14^. The stimulation of methylation is mediated through the recognition of H2Bub by Dot1 and the MLL complex, the methyltransferases for H3K79 and H3K4, respectively ^15,16^. H2Bub further coordinates DNA damage repair factors as part of double-strand break repair mechanisms ^17,18^ and plays an important role in the maintenance of chromatin structure throughout the cell cycle ^19,20^.

In *S. pombe*, H2Bub is deposited by the histone H2B ubiquitin ligase complex HULC, which consists of the Bre1 homologs Brl1, Brl2, the Rad6 homolog Rhp6, and the Lge1 homolog Shf1 ^14,21^. Rhp6 has been shown to modulate heterochromatin strength, with overexpression weakening heterochromatin at the mating type locus, centromeres and telomeres and HULC deletions enhancing heterochromatin features ^21–23^.

Brl1, Brl2 and other Bre1 homologs are RING E3 ligases that form a complex with the E2 conjugating enzyme Rhp6/Rad6. At the heart of the ligation reaction, the RING domain dimer of the E3 ligase activates the E2 conjugating enzyme to transfer ubiquitin from its active site cysteine to the substrate ^24^. The RING domains of Bre1 interact with the acidic patch and DNA of the nucleosome ^25–28^. However, the genetic dependence of H2Bub on all subunits of HULC suggests that the E3 ligase activity is regulated and guided by the coiled-coil regions of Brl1 and Brl2 and by Shf1 (Fig. 1A) ^14,21,29^. In addition, acetylation of Brl1 on lysine 242 in the coiled-coil region has been shown to contribute to the protection of active transcription and promote H2Bub ^30^.

**Figure 1:**
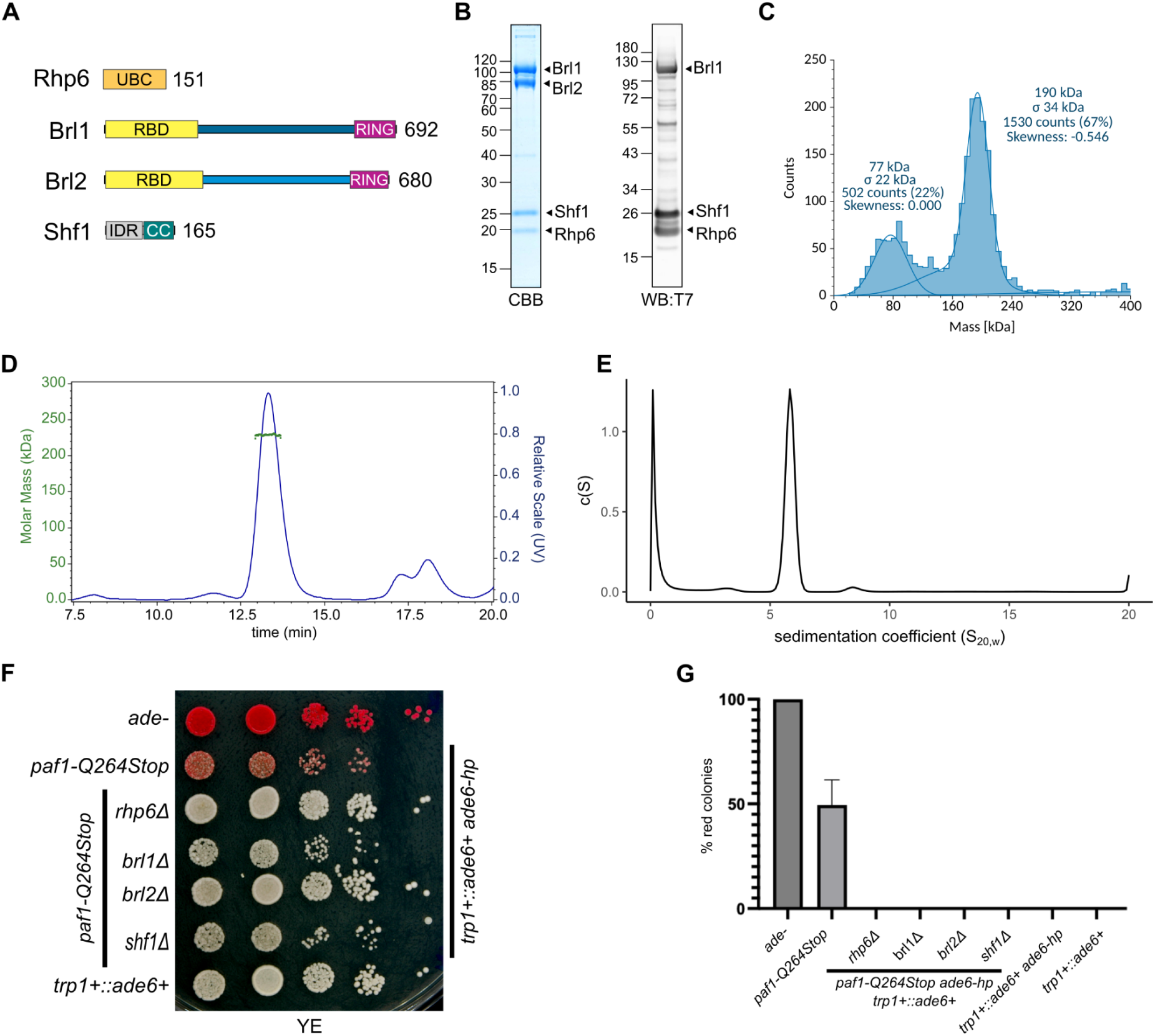
HULC is a Brl1-Brl2-Shf1-Rhp6 assembly with 1:1:1:1 stoichiometry. (A) Domain structure of S. pombe HULC subunits. CC: coiled coil; RING: really interesting new gene domain. RBD: Rad6-binding domain, IDR: intrinsically disordered region, UBC: Ubiquitin C E2 ligase domain (B) SDS PAGE gel of size exclusion peak fraction of HULC and western blot against T7-tagged Brl1, Shf1 and Rhp6 after StrepII-tag pull-down of HULC. (C) Mass photometry size distribution of affinity-purified HULC. (D) Size exclusion coupled with multi-angle light scattering (MALS) showing UV chromatogram overlaid with molecular mass determination for HULC peak. (E) Sedimentation velocity c(S) distribution of HULC after size exclusion chromatography (see fit in Fig. S2). (F) Spot assay of indicated *S. pombe* strains on yeast extract agar (YE). (G) Quantification of red colony frequencies for the mutant strains used in (F). All solution experiments were in 20 mM HEPES pH 7.5, 150 mM KCL, 2 mM MgCl_2_.

In *S. pombe*, small interfering RNAs (siRNAs) function in the initiation and maintenance of heterochromatin ^31–33^. For example, endogenous siRNAs are required to maintain heterochromatin at centromeric repeats. Protein-coding genes, however, are protected from heterochromatin formation even if they are targeted by ectopic siRNAs ^34,35^. *paf1*, *leo1, prf1, mst2*, *dss1*, *mlo3* and *epe1,* which are factors associated with transcription elongation, have been identified as critical for counteracting siRNA-directed heterochromatin formation on protein-coding genes ^36,37^. The Paf1 complex (PAF1C) is part of the transcription elongation apparatus and has been shown to recruit Bre1 to the elongating polymerase in yeast and mammalian systems ^12,38^.

Paf1C consists of five subunits: Paf1, Ctr9, Cdc73, Leo1 and Rtf1, and wraps around the elongating RNA Polymerase II ^39,40^. In *S. cerevisiae*, Rtf1 is part of the Paf1C but its homolog in human and fission yeast shows biochemical and functional separation from the other Paf1C subunits ^41,38,42^. Rtf1 contains a central Plus3 domain, which binds the C-terminal tail of Spt5 that is phosphorylated by Cdk9 (P-TEFb). The Rtf1 N-terminus binds the chromatin remodeler Chd1 and also contains an HMD domain, which recruits the H2B ubiquitin ligase by binding Rad6 ^43^. Furthermore, the HMD domain alone rescues the loss of H2Bub after Rtf1 deletion ^44^. Prf1 is the *S. pombe* homolog of Rtf1 and is part of a feedback loop between Cdk9 and H2B ubiquitination ^42,45^. The Cdk9-Spt5-Prf1-H2Bub axis inhibits transcription elongation and antagonizes a distinct Cdk9-Paf1C pathway, which also impacts H2Bub but promotes transcription elongation.

How HULC and PAF1C interact to control H2Bub remains incompletely understood. Here, we use the *S. pombe* HULC complex as a model system to determine its structural principles and discover that the HMD domain of Prf1 primes HULC for ubiquitination. We determine the stoichiometry of HULC and the structural role of Shf1 in a highly flexible complex. The Prf1HMD domain acts as a scaffold that aligns the RING domains with Rhp6 to activate the ubiquitin transfer to the H2B substrate.

## Results

### 1:1:1:1 Assembly of Brl1-Brl2-Shf1-Rhp6 forms H2B Ubiquitin Ligase Complex

To study the structure and regulation of HULC we co-expressed *S. pombe* Rhp6, Shf1, Brl1 and Brl2 in insect cells and used a tandem StrepII tag on Brl1 to purify the complex (Fig. 1A, Fig. S1). The Coomassie Brilliant Blue (CBB) staining and western blot analysis using a T7 tag engineered into Brl1, Rhp6 and Shf1 subunits shows a roughly equimolar stoichiometry (Fig 1B). However, Rhp6 tends to show weaker intensity compared to Shf1 and Brl1 than expected from a strict one-to-one stoichiometry. Mass photometry revealed molecular mass of 190 kDa (Fig. 1C), and using multi-angle light scattering, we observed a monodisperse peak on gel filtration with a molecular mass of 229 ± 0.5 kDa (Fig. 1D). Analytical ultracentrifugation sedimentation velocity experiments determined a sedimentation coefficient of (S_20,w_) of 6.05S, with an *f/f_0_*frictional ratio of 1.93, corresponding to a heavily elongated protein with a molecular mass of 189 kDa (Fig. 1E, Fig. S2). The calculated mass of HULC with equimolar subunit composition is 218 kDa. Based on the observations in Fig. 1, we conclude that HULC consists of one molecule each of Brl1, Brl2, Shf1 and Rhp6. However, we have observed variability in the intensity of the associated Rhp6, which suggests that it is bound weakly to the complex.

### HULC deletions suppress siRNA-directed silencing in a *paf1-Q264Stop* background

To investigate the role of HULC in heterochromatin silencing, we deleted HULC subunits in an *ade6+* siRNA (ade6-hp) expressing *paf1-Q264Stop* background (Fig. 1F). *paf1-Q264Stop* enables siRNA-directed formation of heterochromatin at the ade6 reporter, which manifests as red colonies ^36^. If H2Bub antagonizes heterochromatin formation as the effect of H2Bub overexpression suggests ^21^, then we would expect siRNA-directed *ade6+* silencing in *paf1-Q264Stop* cells to increase in HULC mutants. However, we observe complete suppression of the silencing by the *paf1-Q264Stop* phenotype upon deletion of every HULC subunit (Fig. 1F,G). This finding is consistent with the data implicating H2Bub in elongation inhibition ^42,45^ and suggests that the *paf1-Q264Stop* mutant affects the stimulation of elongation by PAF1C. By deleting HULC we release the elongation inhibition of the *spt5*-H2Bub axis and thus compensate for the *paf1-Q264Stop* deficiency. These results suggest that transcription elongation rate is a determining factor in siRNA-directed heterochromatin formation on active genes.

### AlphaFold model of HULC predicts modular architecture

Using the 1:1:1:1 stoichiometry, we leveraged AlphaFold (AF) to predict the structure of HULC (Fig. 2A) ^46^. AF consistently produces high-confidence structures of Brl1 and Brl2 assembled into a parallel coiled-coil dimer, which folds back on itself, thereby forming a coiled-coil hairpin (CCH) structure. This brings together the N-terminal RBD and the C-terminal RING domains at the base of the hairpin (Fig. 2B). The Shf1 C-terminus forms part of an α-helical bundle at the center of the CCH, with its IDR protruding from the CCH. HULC can be divided into three structural building blocks consisting of the RBD-Rhp6 complex, the CCH and the RING domains (Fig. 2B). Rhp6 and the RBD form a structural module consistent with the published crystal structure of the Bre1RBD-Rad6 structure ^47^. The RING domains form the canonical dimeric arrangement, which has also been determined for the RING domains of Bre1 ^48^. The RBD is linked to the CCH via Brl1 residues 216-237 and Brl2 residues 202-230, which are likely flexible as judged by their low confidence values. The RING domains are connected to the CCH through shorter but equally flexible linkers (Brl1 626-627 and Brl2 610-614).

**Figure 2:**
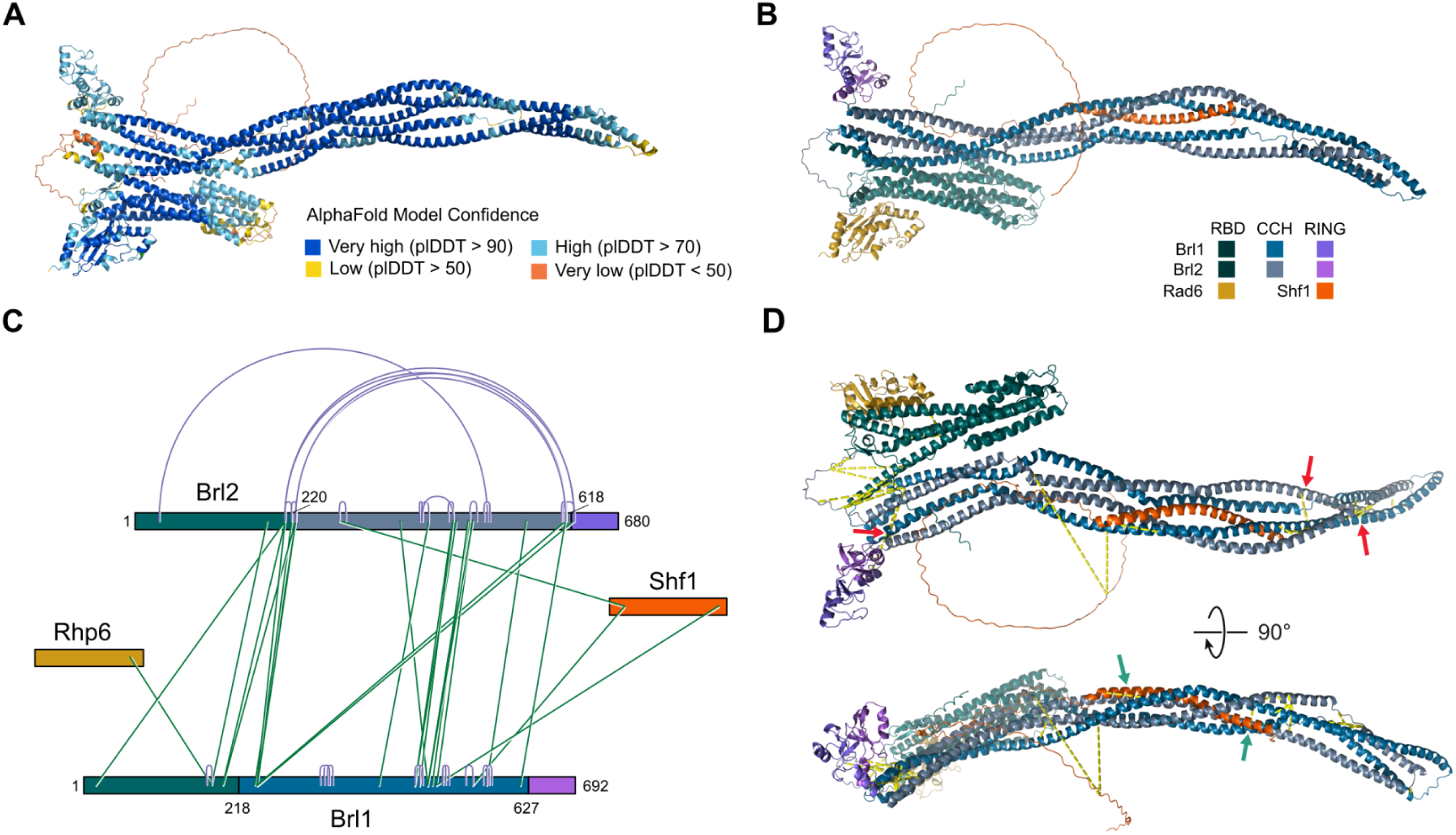
XL-MS supports the AlphaFold-predicted coiled-coil hairpin scaffold of HULC. A) Cartoon representation of the HULC model generated by AlphaFold 3. Colors correspond to pLDDT model confidence scores. B) HULC with Brl1/Brl2 heterodimer colored by structural domains. Shf1 and Rhp6 are colored as in D. C) Schematic representation of contacts within HULC determined by cross-linking mass spectrometry. D) HULC colored by subunit with cross-links supporting the CCH structure highlighted by arrows in red and cross-links consistent with the Shf1-CCH integration highlighted by arrows in teal.

Shf1, corresponding to the IDR-containing subunit of other Bre1-like complexes, has, as expected, a disordered N-terminus (residues 1-191). Its C-terminal α-helix, however, is a fully integrated part of the CCH and ties together the coiled-coils from both prongs of the hairpin. This suggests that it plays a central role in stabilizing the hairpin structure of the CCH.

The interactions between the RBD, CCH and RING domain modules seen in Fig. 2 appear consistently in AF predictions. The RBD nestles its C-terminal Brl1 helix (residues 190-213) into a groove formed by the N-terminal α-helices of Brl1 (238-254) and Brl2 (231-248) at the base of the CCH. The RING domains fold back towards the CCH through polar and ionic interactions in Brl1 (E626-K634, for example). The quaternary arrangement, however, relies on areas of lower model confidence.

To validate the AF model, we employed cross-linking mass spectrometry (XL-MS) using BS3 as a crosslinker (Fig. 2C). The crosslinking pattern we observed is highly consistent with the AlphaFold model. It confirms the parallel arrangement of Brl1 and Brl2 (Fig. 2D, red arrows), and we observe crosslinks that confirm the hairpin structure and its integration of Shf1 (Fig. 2D teal arrows). We also observe a crosslink to Rhp6 that is consistent with the RBD-Rad6 crystal structure and the AF model. Thus, the XL-MS data provides experimental support for the main structural modules of HULC proposed by AF.

### HULC has an elongated, flexible structure

To further validate the AF model and gather information on the shape and flexibility of HULC we subjected HULC to size exclusion chromatography coupled with small-angle X-ray scattering (SEC-SAXS) analysis (Fig. 3A). The dimensionless Kratky plot of HULC shows a curve that rises past the Guinier-Kratky point where globular proteins peak and then stays elevated without returning to the baseline (Fig. 3B). This is indicative of an elongated, flexible molecule. The pair-distance distribution confirms the elongated shape of HULC with an average pair-distance of 119 Å and a maximum pair-distance of 320 Å (Fig. 3C,D). We used DAMMIF to calculate possible sphere models that fit the SAXS data (Fig. 3E). This resulted in metal detector-shaped structures with long handles with a flat mass at their base. Further corroborating the flexible, elongated structure of HULC are the highly heterogenous particles observed when HULC was imaged using negative stain and cryo-electron microscopy (cryo-EM) (Fig. 3F).

**Figure 3:**
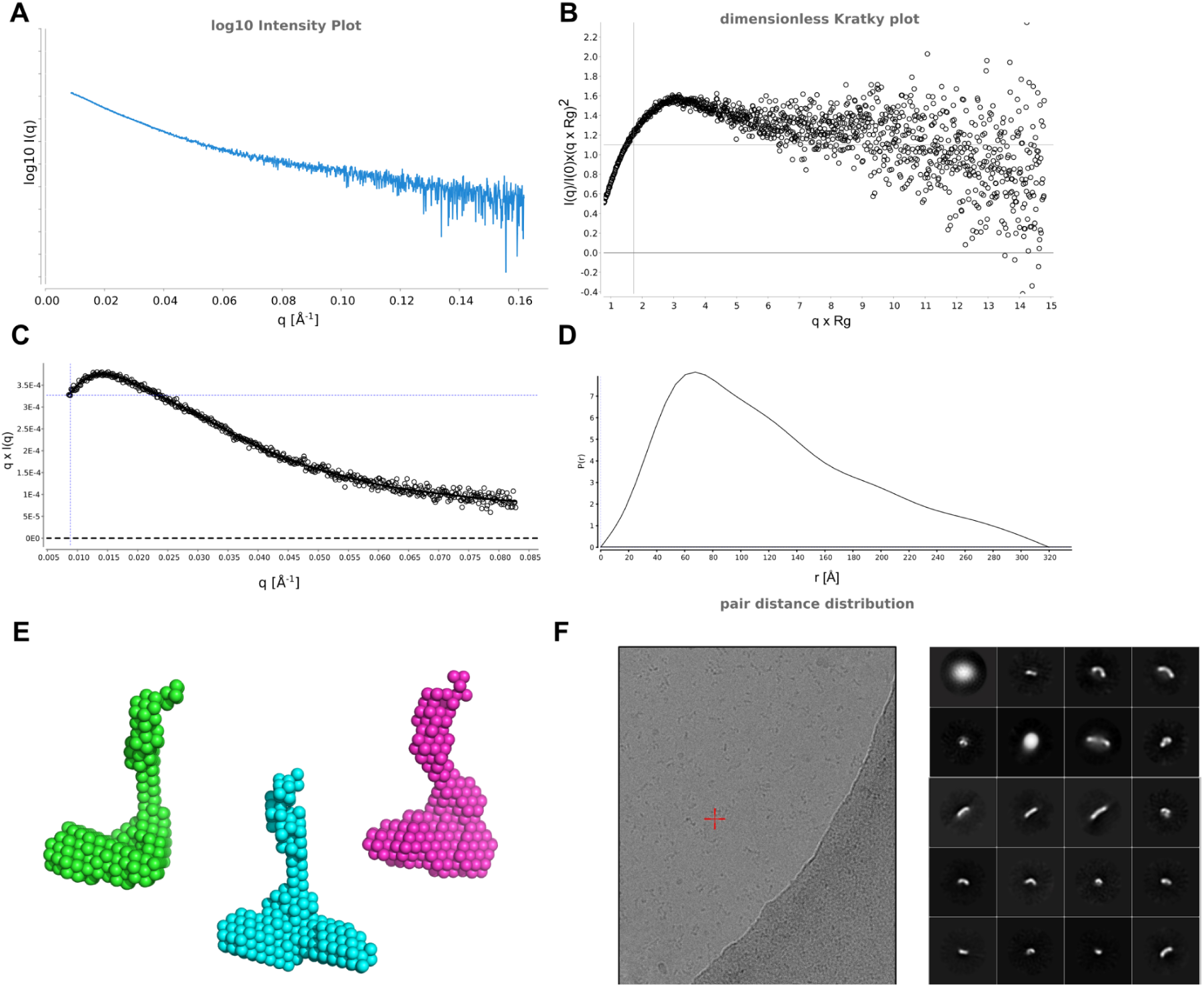
Small-angle X-ray scattering is characteristic of a long, flexible molecule. A) Log_10_ SAXS intensity vs. scattering vector magnitude q of the major SEC peak. Only positive data within the q-range is plotted. B) Dimensionless Kratky plot comparing nucleosome and HULC profiles. Crosshairs indicate the Guinier-Kratky point (1.732, 1.1), the peak position for globular proteins. C) Total scattered intensity plot with pair-distance distribution fit. D) Pair distance distribution function. E) Sphere models were calculated using DAMMIF ^49^. F) Representative cryo-EM micrographs and 2D class averages of HULC particles.

While the SAXS and EM observations are consistent with the elongated structure predicted by AF, they also provide strong evidence that the structure of HULC is heterogeneous and flexible. Based on our structural analysis and the crystal structures of RING and RBD domains ^47,48^, we predict that the individual dimeric modules of HULC - the RBD, CCH and RING domains - form structural entities that are tethered loosely and interact weakly with each other, which leads to high variability in the relative arrangements of these domains. This hypothesis is supported by AF modeling, where we observed that the addition of a tag sequence, for example, can lead to a large rearrangement of the RBD relative to the stalk.

### The HMD domain of the Paf1C subunit Prf1 aligns RING domains and Rhp6 to stimulate HULC activity

Bre1-like complexes are well documented to be recruited to RNA Pol II through Paf1C via multiple contacts between the ubiquitin ligase and different Paf1C subunits ^50,51^. In particular, the HMD domain of Rtf1 has been shown to interact with Rad6 and activate Bre1 ^43,52,44^. We therefore decided to investigate how the HMD-containing N-terminus of Prf1, the *S. pombe* Rtf1 homologue, interacts with the structure of HULC. Intriguingly, the resulting HULC structures showed a large rearrangement of the RBD and RING domains, which was caused by the two structural modules binding side-by-side to the C-terminal ⍺-helix of the HMD domain (Fig. 4A,B, Fig. S3). The binding of Rhp6 is consistent with that proposed for *S. cerevisiae* Rad6 to the Rtf1HMD domain ^43^. The binding of the RING domains to Prf1 unexpectedly involves the N-terminal coiled coil of the Brl1-Brl2 RING domain dimer and an ⍺-helical section within the HMD domain (residues 113-123) that we call the RING-binding Region (RBR) (Fig. 4C).

**Figure 4:**
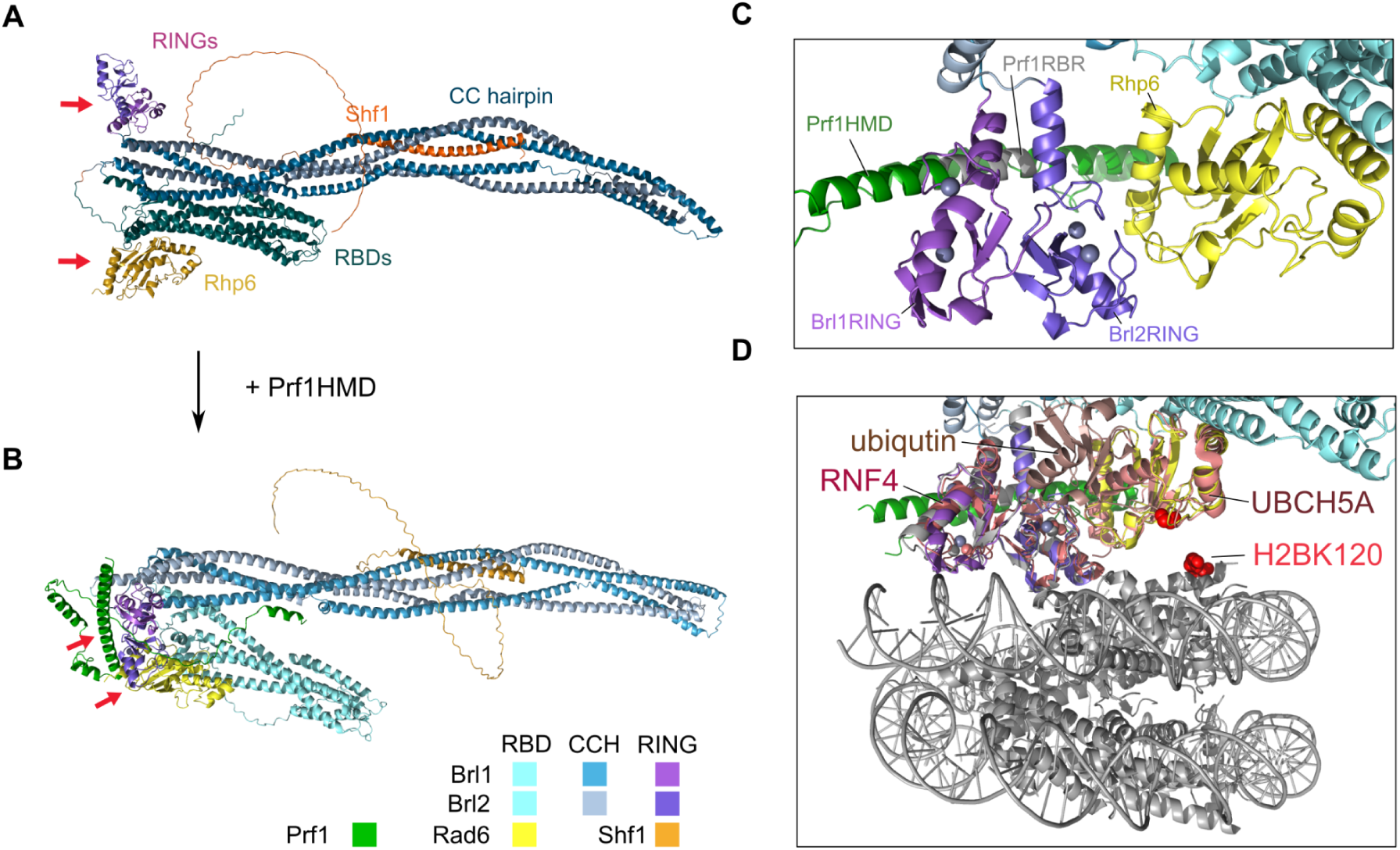
The HMD domain introduces the rearrangement of RBD and RING modules. A) AF model of HULC without the Prf1 N-terminus and B) with the N-terminus of Prf1 (bottom). Arrows indicate the positions of the RING domains and Rhp6 D) Superposition of the HULC-HMD model (C) with the Bre1 RING domains bound to the nucleosome (PDBID:83TY) ^28^ and RNF4-UBCH5A structure (PDBID:4AP4) ^24^. Red colored residues indicate the active site cysteine of Rhp6 and the H2BK120 ubiquitination target.

The catalytic mechanism of RING E3 ligases for the transfer of ubiquitin from the E2 conjugating enzyme to the substrate relies on a specific RING-E2 arrangement that primes the E2-ubiquitin thioester for reaction with the target lysine ^24^. Superposition of the Prf1-bound Brl1-Brl2 RING domains and Rhp6 with the crystal structure of a primed RING-E2-ubiquitin complex (RNF4-UBCH5-Ub, PDBID:4AP4) produces a close 3D alignment with an RMSD of 1.2 Å (Fig. 4D). The ubiquitin is accommodated inside a cradle made up of the HULC RBD, CCH, and RING modules. This is in stark contrast to the AF structure of HULC without Prf1, where the two modules are very far apart from each other (Fig. 4A), and attempts to align them with the RNF4-UBCH5-Ub structure result in RMSDs of around 32 Å. To model substrate binding of Prf1HMD-HULC, we superposed the nucleosome-Bre1RING structures ^26–28^. The resulting primed structure of HULC on the nucleosome shows no clashes, and positions the activated ubiquitin C-terminus at a distance of ∼11 Å from the ε-amino group of the acceptor H2BK120 (Fig. 4D). While not at a sufficiently close distance for catalysis, this proximity would nevertheless be highly conducive for transfer, particularly given the expected flexibility of the system. In conclusion, the AF model of Prf1HMD-HULC suggests that the HMD domain primes the RING domains and Rhp6 for catalysis by orchestrating the alignment for catalysis of the structural modules of HULC.

### Prf1 RBR is required for H2Bub in *S. pombe*

To validate the AF model, we expressed the Prf1 N-terminus of Prf1 comprising the HMD domain in *E. coli* and investigated the effect of Prf1 on HULC activity *in vitro* (Fig. 5A). This data shows that the wild-type Prf1 N-terminus domain stimulates the E3 ligase activity of HULC under our *in vitro* conditions (Fig. 5B). The stimulation depends on the RBR consistent with the AF model and on the concentration of HULC with a pronounced peak at 0.25 µM, where we get a 2-fold stimulation of HULC activity in the presence of the Prf1 N-terminus. This suggests that Prf1 stimulation is critical when the concentrations of HULC are limiting.

**Figure 5:**
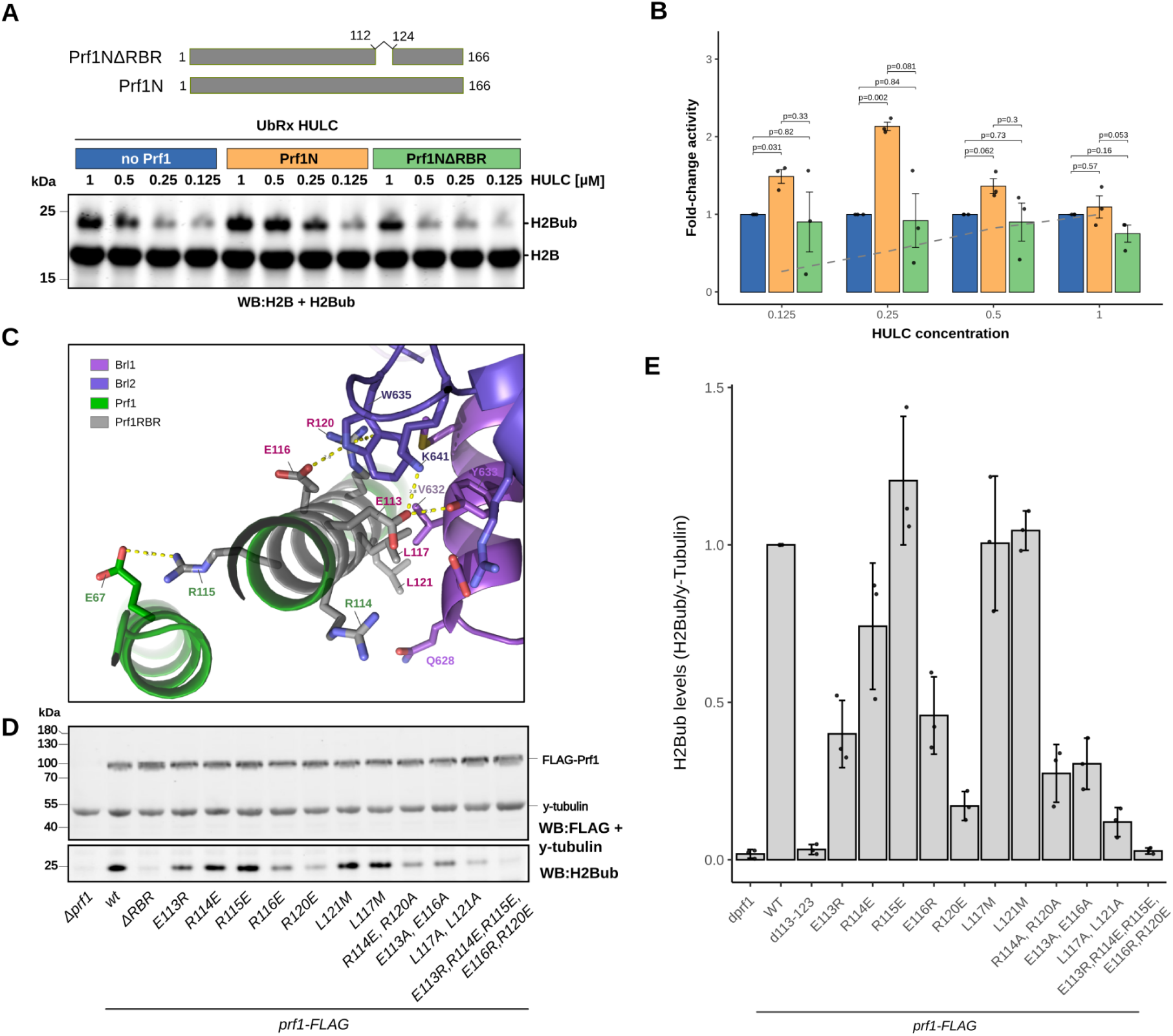
Prf1 modulates the activity of HULC. A) Schematic of the Prf1 constructs used for the *in vitro* ubiquitination assays shown below. Levels of H2Bub and H2B versus differing levels of HULC were determined by Western blotting for the different Prf1 constructs. B) Bar plots overlaid with individual measurements of H2Bub levels normalized to HULC alone. The gray dashed line indicates average activity normalized to 1 µM HULC. Data are presented as mean ± standard error (SE) of three independent experiments (n=3). Statistical significance was determined using paired Student’s t-tests. Fold-change values were calculated by normalizing to the “no Prf1” condition for each HULC concentration. C) Close-up of the RING-Prf1HMD interface predicted by AF. The pink-labeled residues correspond to residues that, when mutated, lose H2Bub compared to wild type. D) Western blot of H2Bub levels in Prf1 mutants. E) Bar plot overlaid with individual measurements of H2Bub levels determined by Western blot and normalized to *prf1-FLAG*.

To investigate the RBR’s role in controlling H2Bub levels *in vivo*, we introduced the RBR deletion as well as point mutations across the RBR into *S. pombe*. The mutations were chosen across the RBR and included single (E113R, E116R, L117M, L121M) and combinations (E113A/E116A, R114A/R120A, L117A/L121A, E113R/R114E/R115E/E116R/R120E) of residues with the potential to disrupt RING binding (Fig. 5C). We also chose to mutate two residues that are close to but not directly involved in the interface (R114E and R115E). Using anti-H2Bub western blotting, we quantified the relative steady-state levels of H2Bub in vegetatively growing *S. pombe* cells (Fig. 5D). A comparison of H2Bub levels shows that the mutants cause a range of effects, with the most severe being indistinguishable from a *prf1* deletion (2% of wild-type levels). The strongest loss is observed in the E113R/R114E/R115E/E116R/R120E (3%) and ΔRBR (3%) mutants. The biggest loss of H2Bub for a single point mutation is observed for R120E (17%), with a similar effect for the R114A/R120A double mutant (27%). E113R and E116R show intermediate loss with H2Bub levels at 40% and 46% of WT and an increased loss for the E113A/E116A combination (30%). L117 and L121 are predicted to form the hydrophobic core of the Prf1-RING interaction, and combined mutations to alanine have a strong effect on H2Bub levels (12%). This can be explained by the loss of considerable hydrophobic bulk and weakening of the hydrophobic core. Interestingly, L117M and L121M mutants have no effect, indicating that increased hydrophobic bulk is easily accommodated. On the other end of the spectrum, with a gain in H2Bub (120%) is R115E. Prf1R115 points away from the Prf1-RING interaction interface and would be expected to have little effect. However, the AF model predicts it to form a salt bridge with the N-terminus of the HMD domain. This could cause a relaxation of the C-terminal ⍺-helix and more optimal alignment of Rhp6 and the RING domains. E113, E116 and R120 cause loss of H2Bub as single and combination mutants. This is consistent with E113 being predicted to form a salt bridge with Brl2K641, E116 forming a hydrogen bond with Brl2W635 and Prf1R120 stacking against the tryptophan side chain of Brl2W635 in a cation-π interaction. The Prf1R114E shows 74% of H2Bub, consistent with being just outside the interaction interface. Thus, the effect we observe on global H2B ubiquitination levels is highly consistent with the prediction that the Prf1RBR aligns RING domains with Rhp6 for ubiquitin transfer to H2B.

### Priming of Bre1-like E3 ligases is also predicted for *S. cerevisiae* and human H2B ubiquitin ligases

Using the insight into the stoichiometry and Prf1HMD-dependent structural changes of *S. pombe* HULC we modeled the Bre1-Lge1-Rad6 complex from *S. cerevisiae* and the RNF20-RNF40-UBE2A-WAC complex from humans (Fig. 6A, C). These models replicate the observed architecture in HULC and share RBD, CCH and RING domain modules. Again, the RBD and RING domains are brought into proximity through the folding back onto themselves of the coiled coils. The species-specific subunits Lge1, WAC and Shf1 all assume a structurally conserved role as integral parts of the CCH.

**Figure 6:**
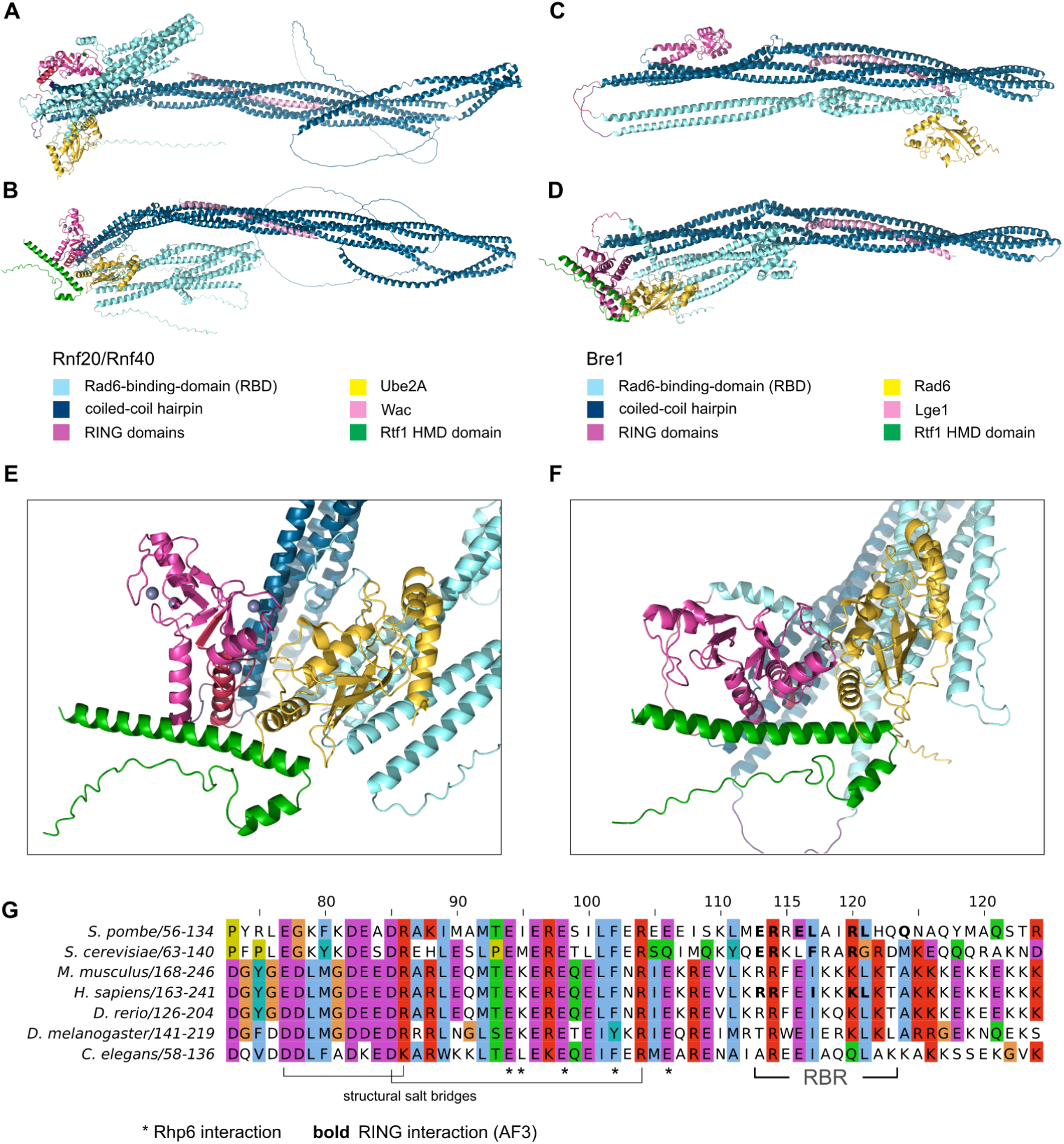
AF models of *S. cerevisiae* and human HULC-HMD complexes predict priming of E3 ligase activity. (A) AF model of human RNF20/RNF40/WAC/UBE2A complex without HMD and (B) with Rtf1 HMD domain. (C) AF model of *S. cerevisiae* HULC complex without HMD and (D) with Rtf1 HMD domain. (E) Close-up of primed state in human HULC. (F) Closeup of primed state in *S. cerevisiae* HULC. (G) Multiple sequence alignment of HMD domains.

We then added the HMD domain of the respective Rtf1 proteins, which, consistent with the observations for HULC, introduced a juxtaposition of the E2 enzyme and the RING domains in a primed configuration (Fig. 6B, D<u>-F</u>). As with HULC, this juxtaposition is associated with major movements of the RING domains and E2 subunit around the base of the CCH. In the case of the *S. cerevisiae* complex, Rad6 localizes near the top of the CCH due to a pocketknife-like opening of the RBD. Adding the Rtf1 HMD domain restores the ⍺-helical bundle of the RBD and arranges Rad6 next to the RING domains. Thus, AF modeling suggests that priming of H2B ubiquitin ligase complexes is conserved across evolution.

Consistent with the structural predictions, we find several conserved residues in the RBR, corresponding to *S. pombe* Prf1 residues R114, E116, L117, R120 and L121 (Fig. 6G)

## Discussion

Our structural and functional analysis of the *S. pombe* HULC complex provides critical insight into the regulatory mechanisms of the highly conserved histone H2B E3 ubiquitin ligase complex HULC. The genetic link between PAF1C and HULC is underscored by the observation that HULC mutants suppress RNAi-induced silencing in Paf1 mutants, suggesting that PAF1C and H2Bub act in a common pathway to maintain an open chromatin state. This functional relationship between PAF1C and HULC prompted us to investigate the molecular basis of how PAF1C controls HULC activity at the structural level.

By combining AlphaFold with biophysical techniques, we obtained a full-length model of a Bre1-type E3 ligase that builds on a dimeric scaffold made of the Bre1-like proteins Brl1 and Brl2. This highly intertwined dimer divides into three structural modules: the Rhp6-binding domain (RBD), a highly elongated coiled-coil hairpin (CCH) and the RING domain dimer.

The experimental structures of the isolated *S. cerevisiae* Bre1 RBD-Rad6 complex and RING domain dimers have been determined previously, and the HULC model is highly consistent with those ^47,48^. The CCH is consistent with the recently published crystal structure of the Bre1-Lge1 CCH ^53^, showing coiled coils of ∼400 amino acids folding back onto themselves and thereby bringing the RING domains and RBD into close spatial proximity despite their wide separation in protein sequence. This provides a structural explanation for how the RBD is able to stimulate the ubiquitin ligase activity ^29,47^. As Shf1 plays an integral role in stabilizing the CCH structure, its loss would abolish the RBD-RING proximity and destabilize the complex, consistent with its critical requirement for H2Bub in *S. pombe* ^14,21^.

Despite the CCH bringing together the RBD and RING domains, the AF model of HULC fails to explain how Rhp6 would interact with the RING domains productively for ubiquitin transfer to the substrate. One of the limitations of AF is the lack of information on the dynamics of protein structures, and it is therefore providing an oversimplified picture of HULC. The SAXS measurements and cryo-EM data reveal significant flexibility in the complex, and we hypothesize that the structured modules of HULC (RBD, CCH and RING domains) dynamically sample a range of relative conformations. It is notable that the *S. cerevisiae* Bre1 complex shows quite drastic structural heterogeneity in the AF models produced, including flipping open of the RBD (Fig. 6D).

Intriguingly, the inclusion of the HMD domain of the Paf1C subunit Prf1/Rtf1 in the AF model leads to the rearrangement of RBD and RING domains into a catalytically conducive arrangement, which we confirm by functional validation in *S. pombe*. The fact that AF predicts very similar structural rearrangements by Rtf1HMD for the *S. cerevisiae* and human complexes strongly suggests that HMD domain-mediated priming is highly conserved across Bre1-type E3 ligases.

The finding that Prf1HMD primes HULC through the RBR-RING interaction raises the question of how this mechanistic insight relates to the genetic observation that HULC mutants suppress RNAi-induced silencing in Paf1 mutants. Since *paf1* mutants lose almost all H2Bub ^30,42,52^, this suppression is unlikely to be mediated through changes in H2Bub levels.

Instead, our results suggest that HULC may contribute to the regulation of transcription elongation through a function that is independent of its ubiquitin ligase activity. In *S. pombe*, Prf1 is an integral part of the Spt5CTD-Prf1 axis, which antagonizes RNA Pol II occupancy independently of PAF1C^42^. The physical recruitment of HULC by Prf1 may therefore contribute to elongation regulation irrespective of H2Bub, and disruption of HULC could tip the balance toward the PAF1C-dependent elongation-promoting axis, suppressing heterochromatin formation. Defining this H2Bub-independent role of HULC will be an important goal for future work.

Our mutagenesis data reveal that sequence changes in the Prf1RBR lead to both loss and gain of H2Bub, showing that the RBR is both critical for H2B ubiquitination and a motif capable of tuning the global levels of H2Bub. H2Bub is important for the regulation of transcriptional programs and the DNA damage response and has been described to have a tumor-suppressive role ^54,55^. Our identification of the RBR as a critical regulatory interface opens up new opportunities to study the interplay between transcription and epigenetics and provides a new target interface for therapeutic interventions in diseases with deregulated H2Bub.

## Materials and methods

### Cloning multi-gene expression constructs

*S. pombe* HULC subunits and a YFP gene were cloned into the pGG1 entry vector using BsmBI and then transferred into pFA6a-based baculo shuttle vectors using BsaI-based GoldenGate cloning (Fig. S1). The pFA6Bac vectors have compatible BsmBI overhangs, which were used to combine the HULC genes into pGGBac, generating pGGBac-HULC.

### Insect cell expression

To generate the recombinant bacmid, the PGGBac-HULC plasmid was transformed into DH10MB bacterial competent cells. The transformed cells were plated on blue-white screening plates containing 50 μg/mL kanamycin, 7 μg/mL gentamicin, 10 μg/mL tetracycline, 100 mg/mL X-gal, and 40 μg/ml IPTG. White colonies were selected, and bacmid DNA was purified by alkaline lysis. Sf9 cells growing in 6-well plates (in ESF921 media, at 27°C) were transfected with purified DNA using FuGENE HD (Promega).

Supernatant (V_0_ virus) was harvested 60 h post-transfection and amplified in two consecutive passages at a multiplicity of infection (MOI) of 0.1 by infecting Sf9 cultured in suspension format. Cell density was maintained between 3 × 10^5^ and 1.5 × 10^6^/mL in ESF921 (Expression Systems). Cell morphology and percentage of YFP positivity were closely monitored. For protein expression, cells at a density of 3 × 10^6^/ml were infected at an MOI of 3 and protein expression was monitored via YFP fluorescence. On >80% of cells showing strong YFP signal (about 60 h post infection), cells were harvested by centrifugation, washed in 1x PBS, flash frozen in liquid nitrogen, and stored at -70°C.

### Protein purification

All steps were performed at 4°C. In a typical preparation, frozen cell pellets were resuspended in purification buffer (20 mM HEPES pH 8, 150 mM KCl, 2 mM MgCl_2,_ 2 mM DTT, 1 mM PMSF, Roche cOmplete EDTA-free Protease Inhibitor Cocktail) to a total volume of 30 ml. Cells were lysed in batches using a Dounce homogenizer with 10–20 strokes with a tight clearance pestle. Lysates were clarified by centrifugation in an SS-34 rotor (Sorvall) at 18,000 rpm for 35 min. Supernatants were filtered (0.45 μm) and applied to a 5 ml StrepTrap HP column (GE Healthcare) pre-equilibrated in binding buffer (20 mM HEPES pH 8, 150 mM KCl, 2 mM MgCl_2,_ 2 mM DTT). The column was washed with 15 column volumes (CV) of wash buffer (20 mM HEPES pH 8, 150 mM KCl, 2mM MgCl_2_ , 2 mM DTT), followed by 5 CV of binding buffer. The protein was eluted using 6 CV of binding buffer supplemented with 5 mM D-desthiobiotin (Merck Life Science Limited).

Elution fractions were concentrated to 500 μL and loaded onto the Superdex 200 Increase 10/300 GL size exclusion chromatography column (GE Healthcare) equilibrated in gel filtration buffer (20 mM HEPES pH 8, 150 mM KCl, 2 mM MgCl_2_). After size exclusion, the protein was concentrated and stored at 4°C.

### Reconstitution of mononucleosomes

Histone octamers and DNA (154 bp Widom 601 DNA) were prepared as described previously ^56,57^. Octamer and DNA were combined at a 0.9:1 ratio in Start buffer (10 mM Tris-HCl, pH 7.5; 1.4 M KCl; 0.1 mM EDTA) and dialyzed into End Buffer (10 mM Tris-HCl pH 7.5, 10 mM KCl, 0.1 M EDTA) at 4°C. Nucleosomes were analyzed on a 5% TBE gel to confirm nucleosome formation.

### Mass photometry

Mass photometry experiments were carried out using a TwoMP mass photometer (Refeyn Ltd., Oxford, UK). Microscope coverslips (630-2187, VWR) and silicone gaskets (GBL103250, Sigma-Aldrich) were cleaned with 50% isopropanol followed by water and dried under a stream of nitrogen. Gaskets were then placed on the cleaned coverslips on the sample stage following the manufacturer’s instructions.

All measurements were performed at least three times independently in separate wells using buffer containing 20 mM HEPES pH 8, 150 mM KCl, and 2 mM MgCl₂. For each experiment, 18 μL of buffer was added to a gasket well, and the focal point was acquired using the autofocus function in Refeyn Acquire MP 2.3.1 software. Then, 2 μL of 200 nM HULC complexes (diluted immediately prior to measurement to prevent complex disassembly) was added to the well and mixed to allow observation of individual landing events below saturation.

Mass photometry movies consisting of 6000 frames were recorded from a 10.8 × 10.8 μm field of view. Data were processed using the default pipeline in Refeyn Discover MP 2.3.0 software. Individual particle contrasts from each video were converted to mass using a contrast-to-mass (C2M) calibration performed in the acquisition buffer. The resulting data were plotted as normalized histograms (bin width = 4.4) and fitted to Gaussian peaks.

For calibration, 2 μL of a 1:100 pre-diluted NativeMark unstained protein standard (LC0725, Thermo Scientific) was added to a gasket well. Masses of 66, 146, 480, and 1048 kDa were used to generate the standard calibration curve in Discover MP software.

### Analytical ultracentrifugation

Sedimentation velocity data were collected using 1.2 cm pathlength cells equipped with quartz windows in a Beckman-Coulter Optima analytical ultracentrifuge. Samples were exchanged into 20 mM HEPES pH 7.5, 150 mM KCL, 2 mM MgCl_2_ and centrifuged at 40,000 rpm in a Ti.50 rotor at 20 °C. Migration through the cell was monitored at 280 nm. 150 Scans were analysed using c(S) distribution analysis through SEDFIT. Correction values for protein buoyancy and buffer density were calculated using SEDNTERP.

### SAXS

SEC-SAXS experiments were conducted at beamline B21 of the Diamond Light Source synchrotron facility (Oxfordshire, UK). Protein samples at concentrations of 10 mg/mL were loaded onto a Superdex 200 Increase 10/300 GL size-exclusion chromatography column (GE Healthcare) equilibrated in 20 mM HEPES pH 8, 150 mM KCl, and 2 mM MgCl₂ at a flow rate of 0.5 mL/min using an Agilent 1200 HPLC system. The column outlet was fed into the experimental cell, and data were collected at 12.4 keV with a detector distance of 4.014 m, recording 3.0 s frames.

Data processing, including frame subtraction, averaging, and Guinier analysis to determine the radius of gyration (Rg), was performed using ScÅtter 3.0. Pair-distance distribution functions [P(r)] were analyzed using PRIMUS, and ab initio shape modeling was carried out with DAMMIF.

### SEC-MALS

Multi-angle light scattering was determined by in-line measurement of the Superose 6 column elution using an Optilab T-rEX differential Refractive index detector connected to a DAWN HELEOS MALS detector (Wyatt Technology). The mass of the complex was calculated using ASTRA software version 6.1.

### Crosslinking mass spectrometry

100 μg of the complex was crosslinked with 0.06 mM of BS3 (Thermo Fisher Scientific), and the reaction was quenched with a 1 M Tris pH 7.5 solution. Samples were precipitated with the Methanol/Chloroform protocol, and then protein pellets were reconstituted in 100 mM Tris (pH 8.5) and 4% sodium deoxycholate (SDC). The proteins were subjected to proteolysis with 1:50 trypsin overnight at 37°C with constant shaking. Digestion was stopped by adding 1% trifluoroacetic acid (TFA) to a final concentration of 0.5%. Precipitated SDC was removed by centrifugation at 10,000 g for 5 min, and the supernatant containing digested peptides was desalted on an OASIS HLB plate (Waters). Peptides were dried and dissolved in 0.5% TFA before liquid chromatography–tandem mass spectrometry (MS/MS) analysis. A total of 1000 ng of the mixture of tryptic peptides was analyzed using an Ultimate 3000 high-performance liquid chromatography system coupled online to an Eclipse mass spectrometer (Thermo Fisher Scientific). Buffer A consisted of water acidified with 0.1% formic acid, while buffer B was 80% acetonitrile and 20% water with 0.1% formic acid. The peptides were first trapped for 1 min at 30 μl/min with 100% buffer A on a trap (0.3 mm by 5 mm with PepMap C18, 5 μm, 100 Å; Thermo Fisher Scientific); after trapping, the peptides were separated by a 50-cm uPAC NEO column (Thermo Fisher Scientific). The gradient was 3 to 35% B in 48 min at 750 nl/min. Buffer B was then raised to 55% in 4 min and increased to 99% for the cleaning step. Peptides were ionized using a spray voltage of 2 kV and a capillary heated at 275°C. The sample was first run using a classic DDA method: the mass spectrometer was set to acquire full-scan MS spectra (350 to 1400 m/z) for a maximum injection time set to Auto at a mass resolution of 120,000 and an automated gain control (AGC) target value of 100%. For 2 seconds, the most intense precursor ions were selected for MS/MS. HCD fragmentation was performed in the HCD cell, with the readout in the Orbitrap mass analyzer at a resolution of 30,000 (isolation window of 1.6 Th) and an AGC target value of 200% with a maximum injection time set to Auto and a normalized collision energy of 30%. Then, it was injected again using a method specific for XL proteomics: the MS1 settings remained the same, but only ions with a charge higher than 3+ were fragmented, and the isolation window was set to 1.2 Th to obtain only clean MS/MS spectra.

All raw files were analyzed first by FragPipe v23 ^58^ software using the automatic standard parameters. Only the proteins identified were then used in the pLink v3 search ^59^, utilizing the pLink standard setting for BS3 XL with an FDR of 1%. Data analysis was then carried out by using PyMol for measuring distances and visualization of crosslinks.

### AlphaFold modeling

HULC models were generated using AlphaFold v3 ^46^. Input sequences contained one each of Brl1, Brl2, Shf1, and Rhp6 with the optional addition of the Prf1 N-terminal domain (1-166).

Four zinc ions were added to complete the RING domains. Models were visualized and inspected in PyMol v3.0 ^60^. Side chain rotamers in the RBR-RING interface were optimized to avoid clashes using Coot ^61^.

### Strain generation

*S. pombe* strains were generated using a PCR-based method ^62^. The *prf1* 5′ UTR and coding sequence were amplified from genomic DNA and cloned into the SmaI site of pMB0364, introducing a C-terminal 3×FLAG tag. Site-directed mutations were introduced into this construct and verified by Sanger sequencing. PCR fragments containing the *prf1* 5′ UTR, coding region, 3×FLAG tag, and hphMX4 cassette—with a primer-derived *prf1* 3′ UTR overhang—were used for transformation of a *prf1Δ* strain. Correct integration was confirmed by colony PCR and protein expression by Western blotting. Strains and plasmids are listed in Tables S1 and S2, and primers in Table S3.

Strains carrying deletions of individual HULC subunits were generated by transforming the reporter strain with a PCR-amplified DNA fragment, as described by Bahler et al. (1998). The fragment contained a *URA3* gene (from *C. albicans*) flanked by homologous sequences upstream and downstream of the target HULC ORF to facilitate homologous recombination. Transformants were selected on appropriate selective media, and the correct integration at the insertion locus was verified by PCR analysis.

### Western blotting

For all the Western blots, samples were separated through SDS-PAGE and transferred to nitrocellulose membrane (Bio-Rad), and the total protein was taken through reactive brown staining. The membranes were blocked overnight at 4°C with LI-COR Intercept blocking buffer (TBS) and immunoblotted with the relevant primary antibody at room temperature for 1 h before being washed. This was followed by incubation with the secondary IRDye antibody and scanning with the Odyssey Imaging System (LI-COR). The images were quantified with the ImageJ software.

| Antibody | Source | Catalogue Number |
| --- | --- | --- |
| Anti-ubiquitinyl Histone H2BK120 monoclonal rabbit antibody | Cell Signalling Technology | 5546S |
| Histone H2B antibody | Active Motif | 39237 |
| Anti-FLAG M2 monoclonal mouse antibody | Sigma Aldrich | F1804 |
| Anti-γ-tubulin monoclonal mouse antibody | Sigma Aldrich | T6557 |
| IRDye 680 anti-mouse IgG secondary antibody | LI-COR | 926-68072 |
| IRDye 800 anti-rabbit IgG secondary antibody | LI-COR | 925-32211 |

### Ubiquitination assay

For the HULC ubiquitination reactions, 100 nM E1 (Ube1, BostonBiochem, Cat# E-304-050), HULC complex, and 30 mM ATP were incubated for 60 minutes with 2 µM mononucleosomes, 1 µM Prf1, and 10 µM ubiquitin at 30°C in reaction buffer (50 mM Tris-HCl, pH 8.0; 50 mM NaCl; 50 mM KCl; and 10 mM MgCl_2_). Reactions were quenched by adding EDTA, pH 8.0, at a final concentration of 50 mM and placing the mixture on ice for 5 min. The reactions were stopped by the addition of a 5X loading buffer. To test the signals of H2B-K120ub, 10 µL of each sample was loaded and analyzed by SDS-PAGE followed by Western blotting with anti-H2B-K120ub antibody and anti-H2B antibody. The signals were quantified using ImageJ.

### Cryo-EM sample preparation and data collection

A 3 µL aliquot of the HULC complex (0.026 mg/mL, ∼0.12 µM) was applied to glow-discharged QUANTIFOIL holey carbon grids (R 1.2/1.3, Cu 300 mesh). Cryo-EM grids were prepared using Vitrobot Mark 4 (Thermo Fisher Scientific) with 100% humidity, ashless filter paper (Standard Vitrobot Filter Paper, Ø55/20mm, Grade 595) and blotting time of 3 sec.

The datasets were collected on a Krios electron microscope operated at 300 kV using a K3 detector (Gatan Inc.) with EPU data acquisition software (ThermoFisher Scientific).

Individual exposures were acquired at a dose rate of 16.9 e⁻ px⁻¹ s⁻¹ in 50 equal fractions over 2.05 s, with a calibrated pixel size of 0.835 Å px⁻¹ and an accumulated dose of 49.7 e⁻ Å⁻². The nominal defocus was varied between −0.8 µm and −2.3 µm in 0.3-µm steps.

All image processing was carried out on the cryo-EM computational cluster, University of Leicester. The raw movies were processed with RELION43. Movies were motion corrected with MotionCor2 and defocus values were calculated with CTFFIND4. Particles were picked using TOPAZ and classified in 2D.

## Acknowledgements

We thank Yvan Pfister for technical support. We thank Diamond Light Source for beamtime (proposal mx34438) and the staff of beamlines B21 for assistance with SAXS data collection. Likewise, we thank Sarah Northall, Komal Ghauri, and Luke Bailey for the generation of preliminary data. We acknowledge the Midlands Regional CryoEM Facility at the Leicester Institute of Structural and Chemical Biology (LISCB), major funding from MRC (MC_PC_17136). Furthermore, we acknowledge UKRI funding support (BB/S018549/1, BB/Y002857/1, BB/Z515826/1) and support from the College of Agriculture and Life Sciences at Cornell University. Research in the lab of M.B is supported by the Novartis Research Foundation.

## Author contributions

**Ammarah Tariq**: Investigation, Writing - Review & Editing; **Shin Ohsawa**: Methodology, Investigation, Writing - Review & Editing; **Riccardo Zenezini Chiozzi**: Investigation, Writing - Review & Editing; **Panagiotis Patsis**: Methodology, Investigation, Writing - Review & Editing; **Charlie Williams**: Investigation, Writing - Review & Editing; **Alessandro Stirpe**: Methodology, Writing - Review & Editing; **Thomas A. Clarke**: Investigation; **Konstantinos Thalassinos**: Resources, **Marc Bühler**: Conceptualization, Writing - Review & Editing; **Thomas Schalch**: Conceptualization, Methodology, Writing - Original Draft, Review & Editing, Funding acquisition.

For the purpose of open access, the author has applied a Creative Commons Attribution license (CC BY) to any Author Accepted Manuscript version arising from this submission

