## Supplemental Information for "The HMD domain of the PAF complex primes Rad6-Bre1 E3 ligase complexes for H2B ubiquitination"

Supplemental Figures

Supplemental Tables

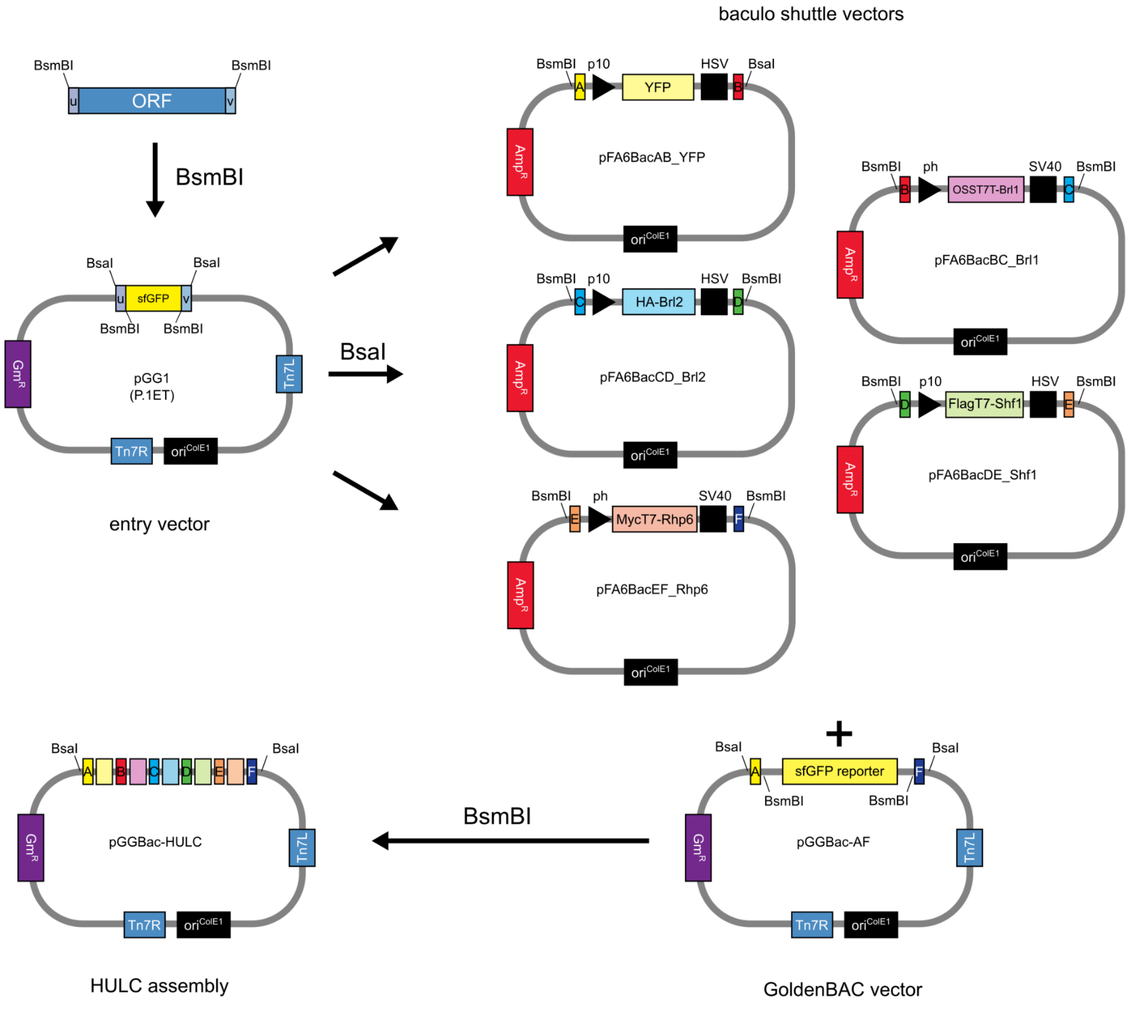


**Figure S1: GoldenGate-based cloning system for HULC expression in insect cells.** The schematic shows plasmids and cloning steps from the pGG1 entry vector to the pGGBAC-HULC baculovirus construct.

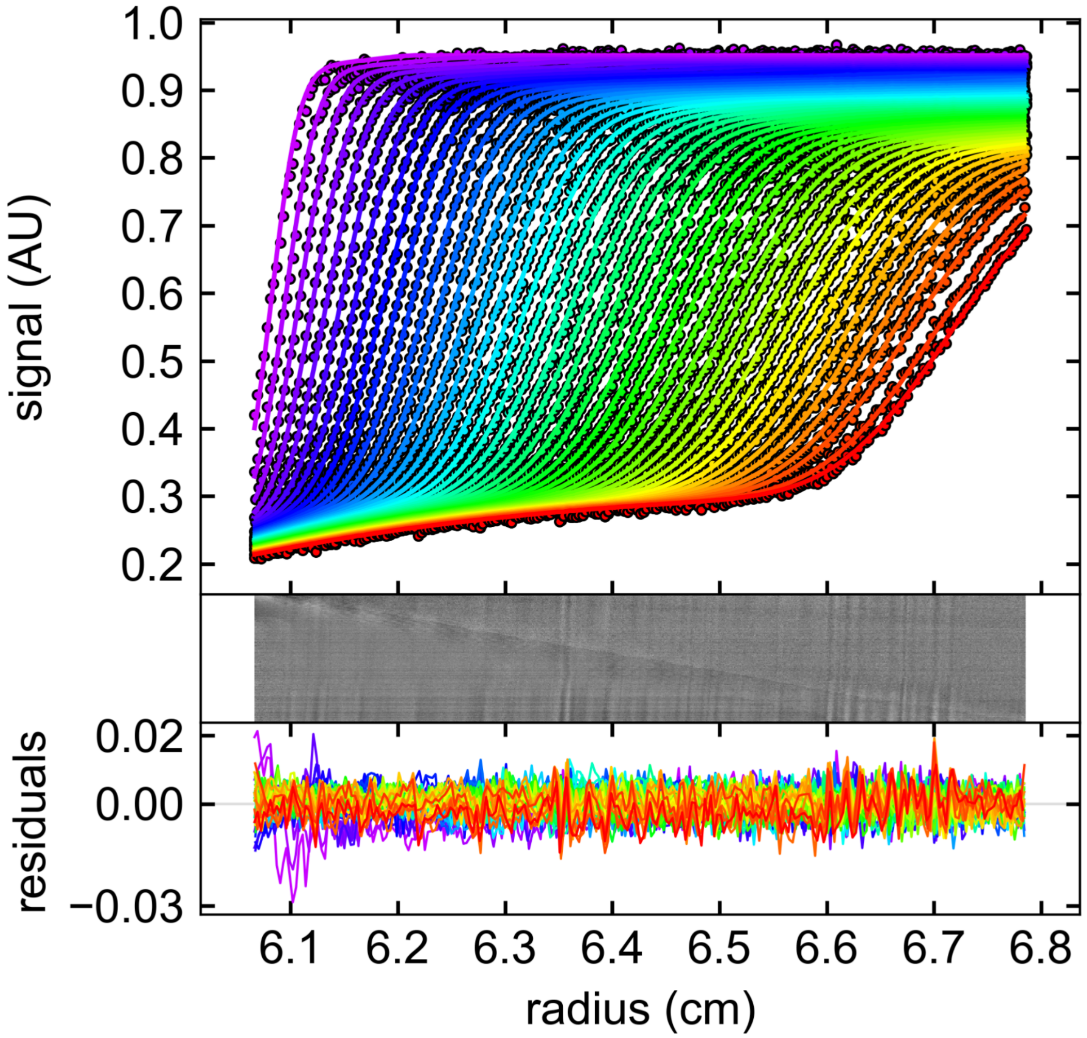


**Figure S2: Analytical ultracentrifugation analysis using Sedfit.** From top to bottom: 280 nm absorbance scans overlaid with fitted data, residual bitmap, and residual plot.

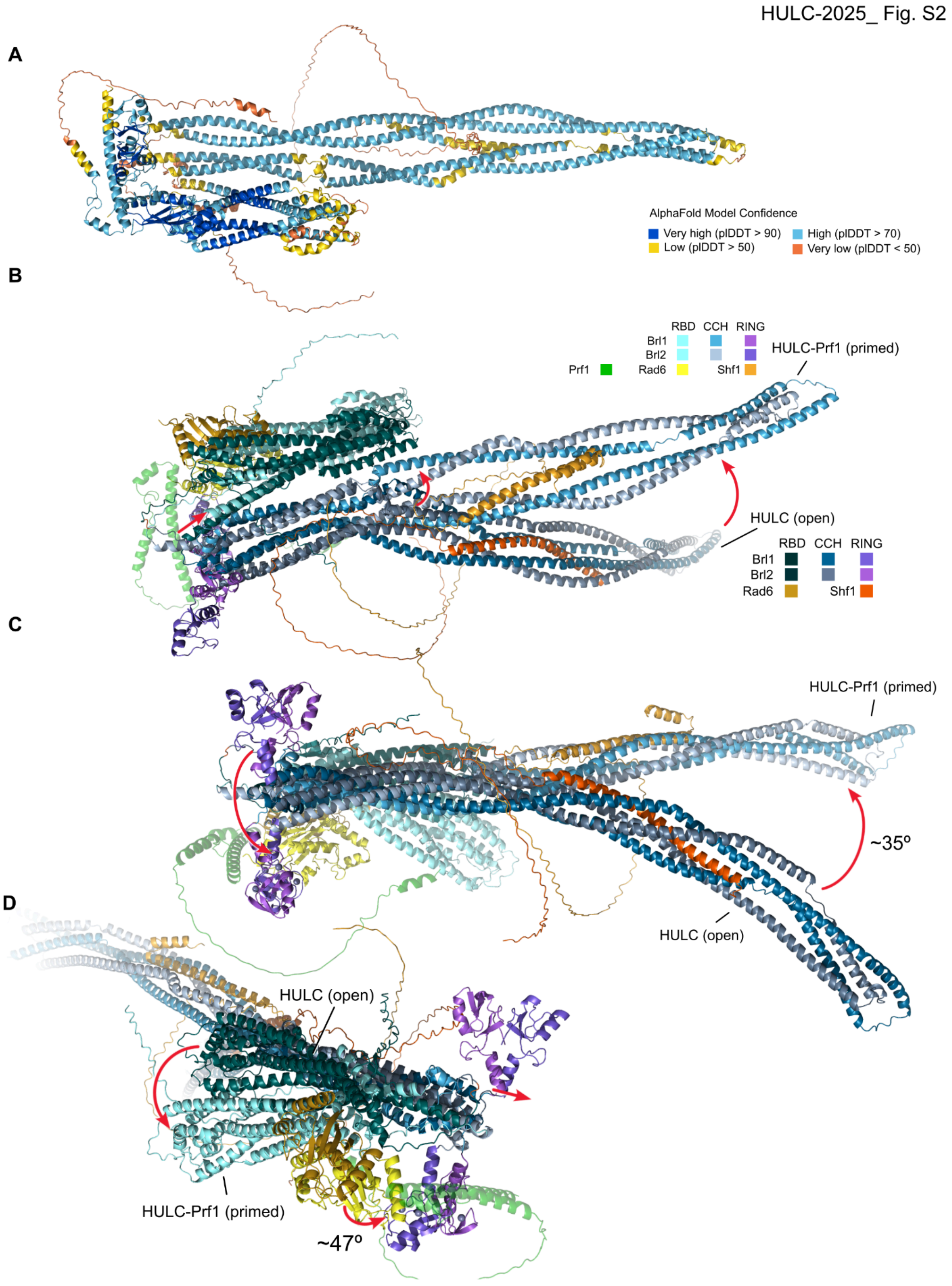


**Figure S3: Structural transition induced by Prf1HMD.** (A) HULC-Prf1 AF3 model colored by model confidence. (B-D) Different views of HULC and HULC-Prf1 superimposed on residues 240-280 of Brl1 show the straightening of the CCH, shift and rotation of the RBD and the flipping of the RING domains.

HULC-2025_ Fig. S4

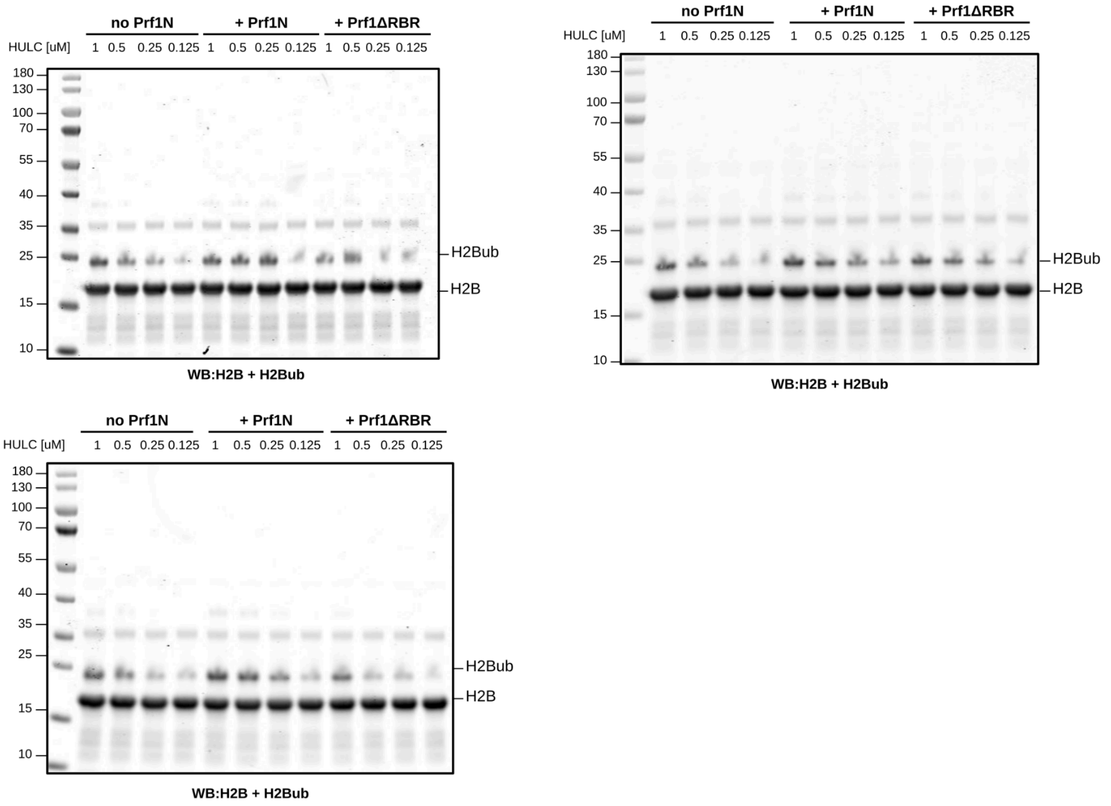


**Figure S4: Western Blots for Prf1 RBR Mutants**. Full Western blot data used for Fig. 5A and B.

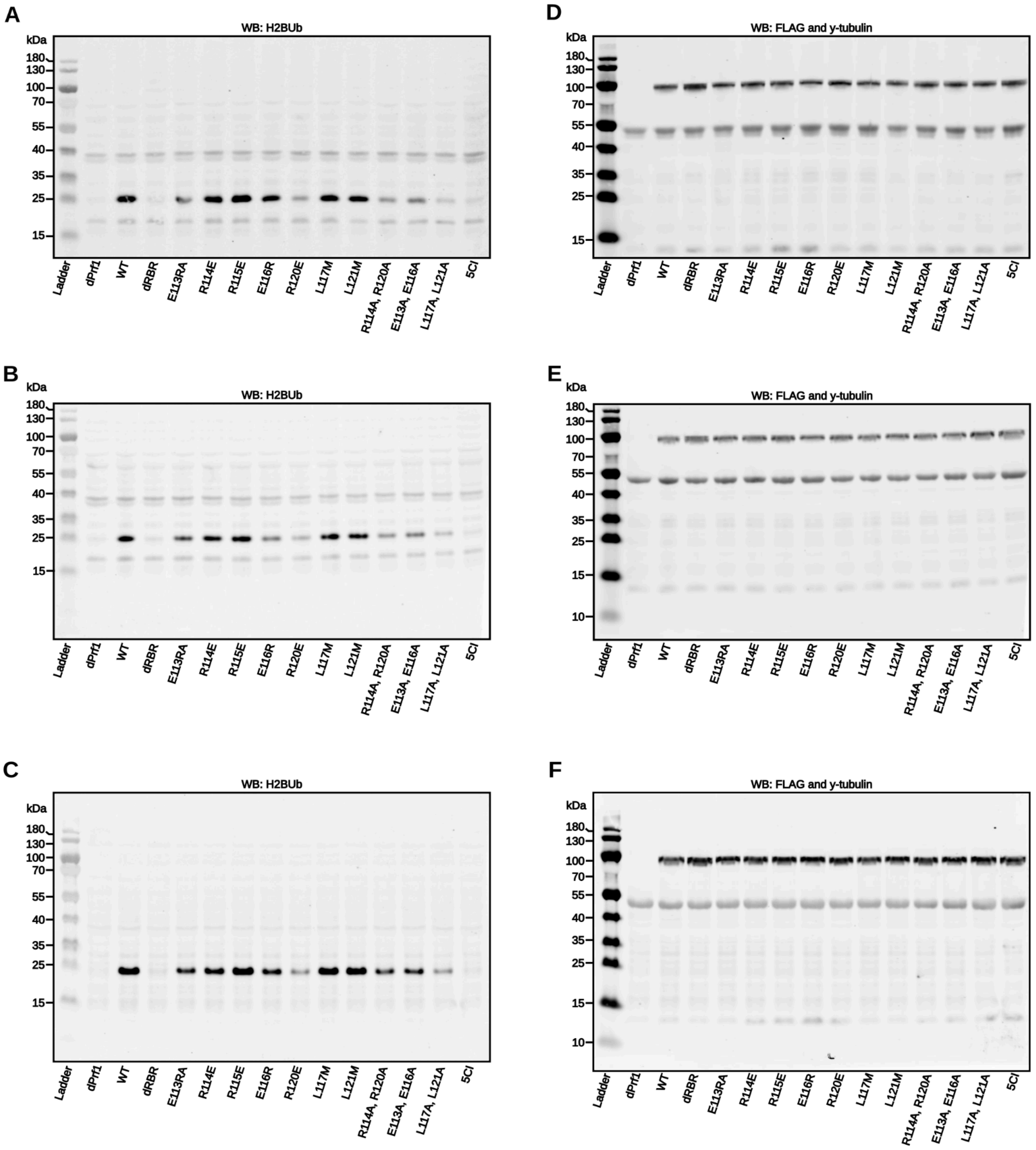


**Figure S5: Western Blots for Prf1 RBR Mutants**. (A-C) Anti-H2BUb and (D-F) anti-FLAG-Prf1 and y-tubulin Western blots used for Fig. 5D and E.

### Supplemental Tables

##### Table S1. Strains used in this study

| Name | Genotype | Figure | Source |
|---|---|---|---|
| SPB2047 | h- leu1-32 ura4-D18 ade6-704 trp1+::ade6+ nmt1+::ade6-hp+::natMX paf1-sms8::LEU2 | 1 F,G | this study |
| SPB4754 | h+ ade6-M216 ura4-D18 leu1-32 prf1Δ::kan | 5 C,E | this study |
| SPB4999 | h+ ade6-M216 ura4-D18 leu1-32 Prf1-6xGly-3xFLAG::hphMX | 5 C,E | this study |
| SPB5300 | h+ ade6-M216 ura4-D18 leu1-32 prf1(113-123d)-6xGly-3xFLAG::hphMX | 5 C,E | this study |
| SPB5314 | h+ ade6-M216 ura4-D18 leu1-32 prf1(E113R)-6xGly-3xFLAG::hphMX | 5 C,E | this study |
| SPB5315 | h+ ade6-M216 ura4-D18 leu1-32 prf1(R114E)-6xGly-3xFLAG::hphMX | 5 C,E | this study |
| SPB5316 | h+ ade6-M216 ura4-D18 leu1-32 prf1(R115E)-6xGly-3xFLAG::hphMX | 5 C,E | this study |
| SPB5317 | h+ ade6-M216 ura4-D18 leu1-32 prf1(E116R)-6xGly-3xFLAG::hphMX | 5 C,E | this study |
| SPB5318 | h+ ade6-M216 ura4-D18 leu1-32 prf1(R120E)-6xGly-3xFLAG::hphMX | 5 C,E | this study |
| SPB5319 | h+ ade6-M216 ura4-D18 leu1-32 prf1(L117M)-6xGly-3xFLAG::hphMX | 5 C,E | this study |
| SPB5320 | h+ ade6-M216 ura4-D18 leu1-32 prf1(L121M)-6xGly-3xFLAG::hphMX | 5 C,E | this study |
| SPB5321 | h+ ade6-M216 ura4-D18 leu1-32 prf1(R114A, R120A)-6xGly-3xFLAG::hphMX | 5 C,E | this study |
| SPB5322 | h+ ade6-M216 ura4-D18 leu1-32 prf1(E113A, E116A)-6xGly-3xFLAG::hphMX | 5 C,E | this study |
| SPB5323 | h+ ade6-M216 ura4-D18 leu1-32 prf1(L117A, L121A)-6xGly-3xFLAG::hphMX | 5 C,E | this study |
| SPB5324 | h+ ade6-M216 ura4-D18 leu1-32 prf1(E113R, R114E, R115E, E116R, R120E)-6xGly-3xFLAG::hphMX | 5 C,E | this study |
| S.0RR | h- leu1-32 ura4-D18 ade6-704 trp1+::ade6+ nmt1+::ade6-hp+::natMX paf1-sms8::LEU2 Brl1::ura3 | 1 F,G | this study |
| S.0RS | h- leu1-32 ura4-D18 ade6-704 trp1+::ade6+ nmt1+::ade6-hp+::natMX paf1-sms8::LEU2 Brl2::ura3 | 1 F,G | this study |
| S.0RT | h- leu1-32 ura4-D18 ade6-704 trp1+::ade6+ nmt1+::ade6-hp+::natMX paf1-sms8::LEU2 Rhp6::ura3 | 1 F,G | this study |
| S.0RQ | h- leu1-32 ura4-D18 ade6-704 trp1+::ade6+ nmt1+::ade6-hp+::natMX paf1-sms8::LEU2 Shf11::ura3 | 1 F,G | this study |

#### Table S2. Plasmids used in this study

| Name | Description | Backbone | Purpose |
|---|---|---|---|
| pMB0364 | pFA6a - 6xGLY - 3xFLAG - hphMX for PCR-based C-terminal FLAG tagging | pFA6a | For backbone of pMB2371 |
| pMB2371 | prf1-6xGly-3xFLAG-hphMX/pFA6a | pMB0364 | For making yeast strain SPB4999 |
| pMB2552 | prf1(E113R)-6xGly-3xFLAG-hphMX/pFA6a | pMB2371 | For making yeast strain SPB5314 |
| pMB2553 | prf1(R114E)-6xGly-3xFLAG-hphMX/pFA6a | pMB2371 | For making yeast strain SPB5315 |
| pMB2554 | prf1(R115E)-6xGly-3xFLAG-hphMX/pFA6a | pMB2371 | For making yeast strain SPB5316 |
| pMB2555 | prf1(E116R)-6xGly-3xFLAG-hphMX/pFA6a | pMB2371 | For making yeast strain SPB5317 |
| pMB2556 | prf1(R120E)-6xGly-3xFLAG-hphMX/pFA6a | pMB2371 | For making yeast strain SPB5318 |
| pMB2557 | prf1(L117M)-6xGly-3xFLAG-hphMX/pFA6a | pMB2371 | For making yeast strain SPB5319 |
| pMB2558 | prf1(L121M)-6xGly-3xFLAG-hphMX/pFA6a | pMB2371 | For making yeast strain SPB5320 |
| pMB2559 | prf1(R114A, R120A)-6xGly-3xFLAG-hphMX/pFA6a | pMB2371 | For making yeast strain SPB5321 |
| pMB2560 | prf1(E113A, E116A)-6xGly-3xFLAG-hphMX/pFA6a | pMB2371 | For making yeast strain SPB5322 |
| pMB2561 | prf1(L117A, L121A)-6xGly-3xFLAG-hphMX/pFA6a | pMB2371 | For making yeast strain SPB5323 |
| pMB2562 | prf1(E113R,R114E,R115E,E116R,R120E)-6xGly-3xFLAG-hphMX/pFA6a | pMB2371 | For making yeast strain SPB5324 |
| pMB2551 | prf1(ΔRBR)-6xGly-3xFLAG-hphMX/pFA6a | pMB2371 | For making yeast strain SPB5300 |
| pGG1 (P.1ET) | Entry vector for schalch lab Goldengate system | | For making GoldenBAC vector containing HULC complex subunits |
| pGGBac-AF | GoldenBAC vector AF containing sfGFP reporter to aid with cloning. Accepts entry vectors AB-EF. | | For making GoldenBAC vector containing HULC complex subunits |
| pFA6BacAB_YFP | GoldenBAC entry vector AB containing YFP with p10 promoter and HSV terminator. | MultiBac | For making GoldenBAC vector containing HULC complex subunits |
| pFA6BacBC_OSST7T-Blr1 | GoldenBAC entry vector BC containing Strep-sumoStar tagged Brl1 with polyhedrin promoter and SV40 terminator. | | For making GoldenBAC vector containing HULC complex subunits |
| pFA6BacCD_HA-Brl2 | GoldenBAC entry vector CD containing HA-tagged Brl2 (no T7 tag) with p10 promoter and HSV terminator. | | For making GoldenBAC vector containing HULC complex subunit |
| pFA6BacDE_FlagT7-Shf1 | GoldenBAC entry vector DE containing Flag-T7 tagged Shf1 with p10 and HSV terminator. | | For making GoldenBAC vector containing HULC complex subunit |
| pFA6BacEF_MycT7-sfotp-Rhp6 | GoldenBAC entry vector EF containing Myc-T7 tagged Rhp6 with polyhedrin promoter and SV40 terminator. Codon optimized for *S. frugiperda*. | | For making GoldenBAC vector containing HULC complex subunit |

**Table S3. Primers used in this study**

| Name | Description | Backbone | Purpose |
|---|---|---|---|
| mb18335 | ggtcgacggatccccgggATCAACGCA GATCCGTTTTC | genomic DNA | Forward primer for cloning of prf1 with mb18336. Lower-case letters indicate overhang sequeince to insert into pMB0364 SmaI site. |
| mb18336 | cccgttaattaacccgggAATGTTGATAT CAATGCCAAAATC | genomic DNA | Reverse primer for cloning of prf1 with mb18335. Lower-case letters indicate overhang sequeince to insert into pMB0364 SmaI site. |
| mb19816 | TTGATGAGGAGGCGTGAATTG GCAATT | pMB2371 | Forward primer for site directed mutagenesis of E113R of prf1 with mb19817. Lower-case letters indicate mutation. |
| mb19817 | ACGCCTCCTCATCAACTTTGAA ATTTC | pMB2371 | Reverse primer for site directed mutagenesis of E113R of prf1 with mb19816. Lower-case letters indicate mutation. |
| mb19818 | ATGGAAgaaCGTGAATTGGCAAT TCGC | pMB2371 | Forward primer for site directed mutagenesis of R114E of prf1 with mb19819. Lower-case letters indicate mutation. |
| mb19819 | TTCACGttcTTCCATCAACTTTGA AAT | pMB2371 | Reverse primer for site directed mutagenesis of R114E of prf1 with mb19818. Lower-case letters indicate mutation. |
| mb19820 | GAAAGGgaaGAATTGGCAATTC GCCTC | pMB2371 | Forward primer for site directed mutagenesis of R115E of prf1 with mb19821. Lower-case letters indicate mutation. |
| mb19821 | CAATTCttcCCTTTCCATCAACTT TGA | pMB2371 | Reverse primer for site directed mutagenesis of R115E of prf1 with mb19820. Lower-case letters indicate mutation. |
| mb19822 | AGGCGTaggTTGGCAATTCGCCT CCAT | pMB2371 | Forward primer for site directed mutagenesis of E116R of prf1 with mb19823. Lower-case letters indicate mutation. |
| mb19823 | TGCCAAcctACGCCTTTCCATCA ACTT | pMB2371 | Reverse primer for site directed mutagenesis of E116R of prf1 with mb19822. Lower-case letters indicate mutation. |
| mb19824 | GCAATTgaaCTCCATCAGCAAAA TGCT | pMB2371 | Forward primer for site directed mutagenesis of R120E of prf1 with mb19825. Lower-case letters indicate mutation. |
| mb19825 | ATGGAGttcAATTGCCAATTCAC GCCT | pMB2371 | Reverse primer for site directed mutagenesis of R120E of prf1 with mb19824. Lower-case letters indicate mutation. |
| mb19826 | CGTGAAatgGCAATTCGCCTCCA TCAG | pMB2371 | Forward primer for site directed mutagenesis of L117M of prf1 with mb19827. Lower-case letters indicate mutation. |
| mb19827 | AATTGCcatTTCACGCCTTTCCAT CAA | pMB2371 | Reverse primer for site directed mutagenesis of L117M of prf1 with mb19826. Lower-case letters indicate mutation. |
| mb19828 | ATTCGCatgCATCAGCAAAATGC TCAG | pMB2371 | Forward primer for site directed mutagenesis of L121M of prf1 with mb19829. Lower-case |

| | | | |
|---|---|---|---|
| | | | letters indicate mutation. |
| mb19829 | CTGATGcatGCGAATTGCCAATT CACG | pMB2371 | Reverse primer for site directed mutagenesis of L121M of prf1 with mb19828. Lower-case letters indicate mutation. |
| mb19830 | gcaCGTGAATTGGCAATTgcaCTC CATCAGCAAAATGCT | pMB2371 | Forward primer for site directed mutagenesis of R114A, R120A of prf1 with mb19831. Lower-case letters indicate mutation. |
| mb19831 | tgcAATTGCCAATTCACGtgcTTC CATCAACTTTGAAAT | pMB2371 | Reverse primer for site directed mutagenesis of R114A, R120A of prf1 with mb19830. Lower-case letters indicate mutation. |
| mb19832 | TGgcaAGGCGTgcaTTGGCAATTC GCCTCCATCA | pMB2371 | Forward primer for site directed mutagenesis of E113A, E116A of prf1 with mb19833. Lower-case letters indicate mutation. |
| mb19833 | AAtgcACGCCTtgcCATCAACTTT GAAATTTCTT | pMB2371 | Reverse primer for site directed mutagenesis of E113A, E116A of prf1 with mb19832. Lower-case letters indicate mutation. |
| mb19834 | gcaGCAATTCGCgcaCATCAGCAA AATGCTCAG | pMB2371 | Forward primer for site directed mutagenesis of L117A, L121A of prf1 with mb19835. Lower-case letters indicate mutation. |
| mb19835 | tgcGCGAATTGCtgcTTCACGCCT TTCCATCAA | pMB2371 | Reverse primer for site directed mutagenesis of L117A, L121A of prf1 with mb19834. Lower-case letters indicate mutation. |
| mb19836 | agggaagaaaggTTGGCAATTgaaCTC CATCAGCAAAATGCT | pMB2371 | Forward primer for site directed mutagenesis of E113R,R114E,R115E,E116R,R120E of prf1 with mb19837. Lower-case letters indicate mutation. |
| mb19837 | ttcAATTGCCAAcctttcttccctCATCA ACTTTGAAATTTC | pMB2371 | Reverse primer for site directed mutagenesis of E113R,R114E,R115E,E116R,R120E of prf1 with mb19836. Lower-case letters indicate mutation. |
| mb19734 | GCATTTTGCATCAACTTTGAAA TTTCTTC | pMB2371 | Forward primer for site directed mutagenesis for ΔRBR(113-123) of prf1 with mb19735 |
| mb19735 | AGTTGATGCAAAATGCTCAGTA TATGGC | pMB2371 | Reverse primer for site directed mutagenesis for ΔRBR(113-123) of prf1 with mb19734 |
| mb18335 | GGTCGACGGATCCCCGGGATC AACGCAGATCCGTTTTC | All pMB2371 | Forward primer for site directed mutagenesis of L117M of prf1 with mb19827. Lower-case letters indicate mutation. |
| mb14430 | aaaagctgacaggtttatataatcctggatgattttt gcaaactaggataataataaataaaacgcgagaat actgagaAGCGCGTTGGCCGATTC ATTA | | Reverse primer for site directed mutagenesis of L117M of prf1 with mb19826. Lower-case letters indicate mutation. |
